# Enhancement of non-specific immune responses in European sea bass (*Dicentrarchus labrax*, L. 1758) by geophyte extract injections (*Urginea maritima* and *Muscari comosum*)

**DOI:** 10.1101/180851

**Authors:** Gülşen Uluköy, Esin Baba, Ramazan Mammadov

## Abstract

The immunomodulatory effects of bulb extracts from the geophyte plants *Muscari comosum* (MC) and *Urginea maritima* (UM) on a non-spesific immune responses of European sea bass were investigated. Ethanol extracts were administered via intraperitoneal injection at doses of 0.5mg/fish and 2mg/fish in PBS. Non-spesific immune parameters such as Nitro blue tetrazolium (NBT) positive cells, serum lysozyme, total protein, total number of leukocytes, leukocyte cell type percentages and specific growth rates were investigated on the 1st, 7th, 14th, 21st, 28th and 35th days following the injection. The results indicate that receiving injections with MC and UM enhances some non-specific immune parameters in European sea bass. Total protein in serum was not enhanced. Activation occured on the 14^th^ day and 21^st^ day and this positive effects started to decrease following days. The appropriate dosage to achieve this enhancement was determined to be 0.5 mg/fish for UM and 2 mg/fish for MC. This preliminary study indicates that these doses yield the best results to promote the health status of European sea bass in intensive aquaculture.

## 1. Introduction

The European Sea bass *Dicentrarchus labrax* is a saltwater fish, belonging to the family Moronidae. European sea bass is cultured in both extensive and intensive systems in the Mediterranean Sea region. Different studies have reported that natural products can be used as alternative components in the diets of European sea bass. Many biological immunostimulant have been found to be effective in European sea bass (Bagni et al. 2000; Bagni et al. 2005; Torrecillas et al. 2007; Yılmaz and Ergün, 2012; Yılmaz et al. 2012). Common immunostimulants are biocompatible, biodegradable, cost effective and safe for the earth (Ortuno et al. 2002). Up to now an extensive number of conventional medicinal plants have been utilized for control of several infections in fish (Duke 1985).

Two herbs were selected for the current study. *Muscari comosum* (MC) commonly called the tassel grape hyacinth, is a geophyte plant that has known diuretic, anti-inflammatory and hypoglycemic activities and has stimulant compounds in its bulb extract (Villa et al. 2012; Nasrabadi et al. 2013; Loizzo et al. 2010). Also, it has antioxidant activity (Pieroni et al. 2002) which is shown with 1.1.-diphenyl-2-picrylhydrazil (DPPH) radical assay. MC bulbs contain mucilages, sugars, latex, tannins, waxes and traces of votalite oil (Davis and Stuart, 1984). *Urginea maritima* (UM), ordinarily called sea squill or sea onion, also has been used for medicinal purposes. The bulb contains cardiovascular glycosides which encourage the heart in human. The most dynamic compounds in the plant are scillirosides, particularly proscilloridine A (Chase et al. 2009). Different constituents found in sea squill incorporate flavonoids, sinistrin (a carbohydrate similar to inulin) and related carbohydrates (Praznik and Spies, 1993). UM has been utilized generally by botanists, chiefly for its impact upon the heart and for its empowering, expectorant and diuretic properties. The fresh bulb is somewhat more dynamic therapeutically than the dried bulb. UM extract has been reported to exhibit peripheral vasodilation in anesthetized rabbits (Barnes et al., 2007).

The present study aimed at determining the immunostimulating effect of these two geophyte plants on non-specific immune reponses and the specific growth rate in European sea bass.

## 2. Materials and methods

### 2.1. Collecting and Extracting of geophyte plants

The geophyte plants (MC and UM) were gathered from their natural habitat in the Mugla-Turkey. The plants were excavated from ground with their bulbs. The bulbs and plant were thoroughly washed under water to remove debris and the soil. Thereafter, bulbs were removed from plant and peeled. Afterward, bulbs minced into little pieces and air dried. These pieces (200–250 g) were added into 500 mL flasks and topped with ethyl alcohol (70%) up to 500 mL. This suspension was left in a temperature adjusted shaking water bath at 50 °C and stayed there for 24 h. It was then filtered in a Whatman filter (0.45 μm). The plant residue was re-extracted with addition of a new ethyl alcohol (70%) as before and the process was repeated three times. The combined filtrates were concentrated on a rotary evaporator at 45°C for ethanol elimination. Then extracts were lyophilized and kept at 4 °C in dark bottles (Lee et al. 2000).

### 2.2. Fish

European sea bass *Dicentrarchus labrax* (90 ± 5 g), a total 300 fish were obtained from a commercial farm at Yalıkavak/Bodrum, Muğla-Turkey and fish were placed in the same farm in six 500 L tanks for the trial. They were acclimated for 2 weeks. Fish were fed with commercial pellets at 2% body weight and water quality parameters were monitored daily.

### 2.3. Experimental design

The trial groups were formed as three fish groups and each group was duplicated. For each group a total of 50 fish was placed into tanks. Each fish in the experimental groups received 100 μl of the plant extract as 0.5 mg/fish and 2 mg/fish by intraperitonal injection (i.p.). For the injection, obtained extracts were dissolved in sterile phosphate buffered saline solution (PBS) and applied using a 26 G needle and 1mL syringe. The control group received only 100 μl PBS.

### 2.4. Preparation of serum and blood cells

Fish samples were randomly selected from each tank after injection and fish were anesthetized by using phenoxyethanol, on the 1^st^, 7^th^, 14^th^, 21^st^ and 28^th^ days. Then blood samples were taken from the fish caudal vein in two steps. A part of blood (∼ 500 μl) was separated directly into an Eppendorf tube to make up serum for assaying lysozyme and total protein. To be able to obtain the serum from blood, the samples were kept at 4 °C overnight and then centrifuged at 3500 g for 15 minutes. The collected serum samples were stored at – 20 °C until assayed. During the blood sampling a part of the blood (∼ 1000 μl) was taken with a heparinized syringe for other tests and used it immediately.

### 2.5. Respiratory burst activity

The Nitro blue tetrazolium (NBT) glass adherent assay was used for determination of the respiratory burst activity in the blood. The assay was carried out following the protocol of Anderson et al. (1992) with some modifications. For this purpose, sampled The NBT positive cells were defined as the phagocytic cells which showed activation and were seen as dark blue in color under the microscope. For each fish, 25 fields were counted on the slide and then the mean and standard error per field were calculated.

### 2.6. Lysozyme activity

The turbidimetric assay was used to determine lysozyme activity by using a modified method which described by Parry et al. (1965) and Hutchinson and Manning (1996). The absorbance at 490 nm. One unit of lysozyme activity was defined as a reduction in absorbance of 0.001/ min.

### 2.7. Serum total protein

The Bradford reagent (Sigma, B6916) used to determine the concentration of total proteins in fish blood (Bradford 1976). The absorbance was measured at 595 nm. The total protein in serum was calculated by using a standard (Bovine Serum Albumin) curve and expressed as mg/mL.

### 2.8. Hematology

Total leukocyte counts (WBC) were made from the fresh blood samples in each fish by using a Neubauer counting chamber as described by Schaperclaus et al. (1991). The hematocrit value was measured overlaying the hematocrit tubes on a sliding scale hematocrit reader (Steinhagen et al. 1990; Schaperclaus et al. 1991).

### 2.9. Growth

Individually, the weights of fish in each group were measured in the beginning and the end of trial. Specific growth rate (SGR) (Laird and Needham (1988));

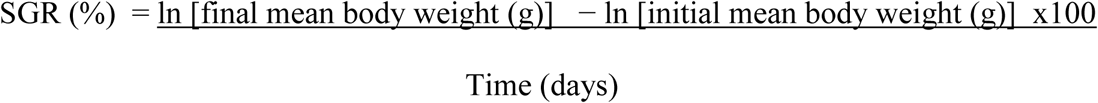

### 2.10. Statistical analysis

Statistical analysis of data involved one-way analysis of variance (ANOVA) followed by Tukey’s multiple pair wise comparison test. The data were expressed as arithmetic means with standard error (SE). Statstical significance was assumed at P < 0.05.

## 3. Results

### 3.1. Respiratory burst activity

The respiratory burst activities are shown Figure 1. The data show that extracellular production of oxidative radicals in fish blood which received 0.5mg/fish UM or MC and 2mg/fish UM or MC experimental groups were higher than the values of the control group (P<0.05). Significantly higher levels of extracellular burst were observed on day 21 in 0.5mg/fish UM and 2mg/fish MC experimental groups (Figure 1). This result shows that application of UM and MC causes an enhacement in the activity of phagocytic cells starting with the first day post-treatment in experimental groups and showed the highest level at the 21^st^ day in 2mg/fish treatment in both groups compare to control.

**Figure 1.**
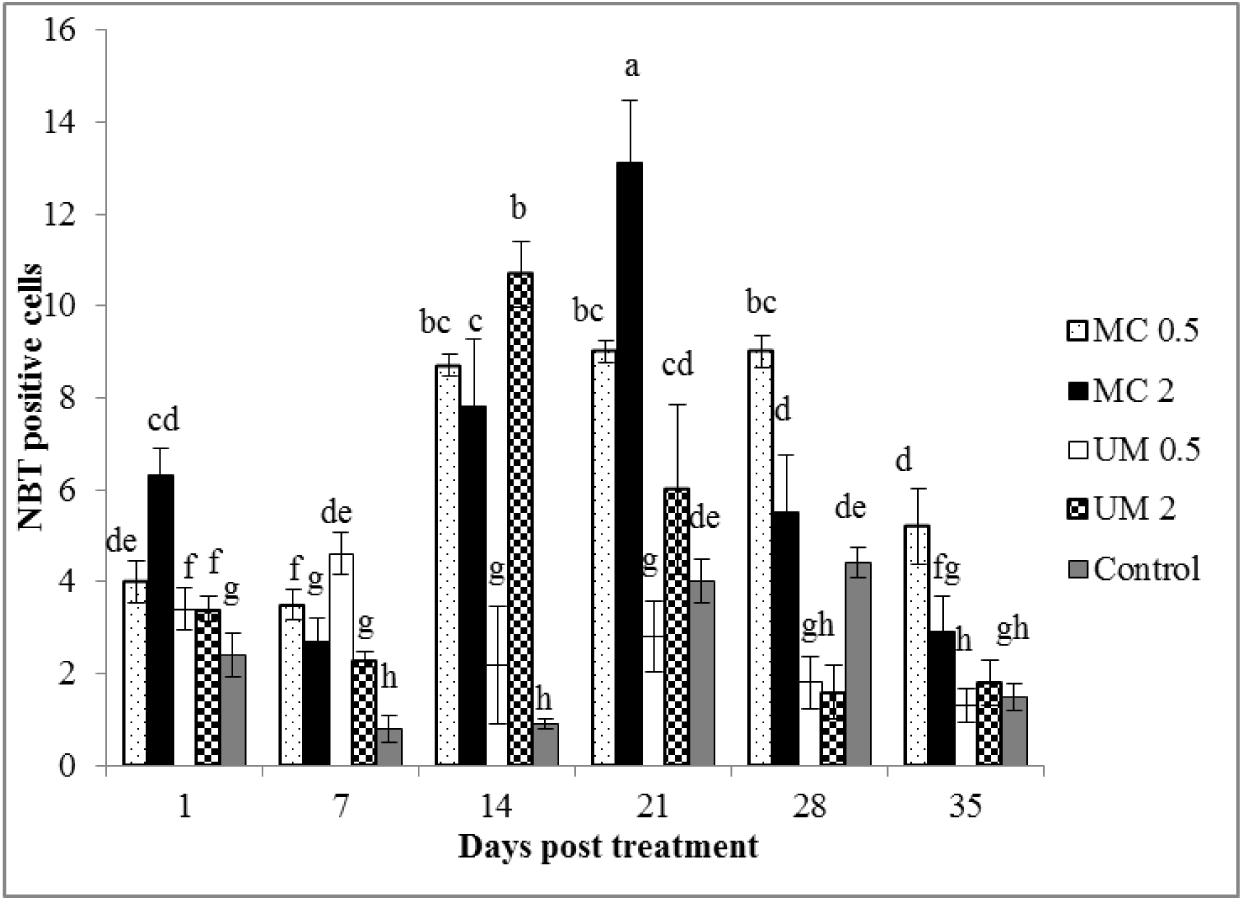
Effects of MC and UM extracts on the number of NBT-positive cells in blood of European sea bass. Data are expressed as means ± SE (n = 10). Mean values at bars with different superscript letters at same stage are significantly different (P < 0.05) from the control.

### 3.2. Serum lysozyme activity

The lysozyme activities in serum are shown in Figure 2. It can be observed that, there are differences among the experimental groups and the control group on certain days (P < 0.05). It has been detected that the the maximum level of lysozyme activities on day 21 in 0.5mg/fish UM and day 7, 14 in 2 mg/fish MC experimental groups.

**Figure 2.**
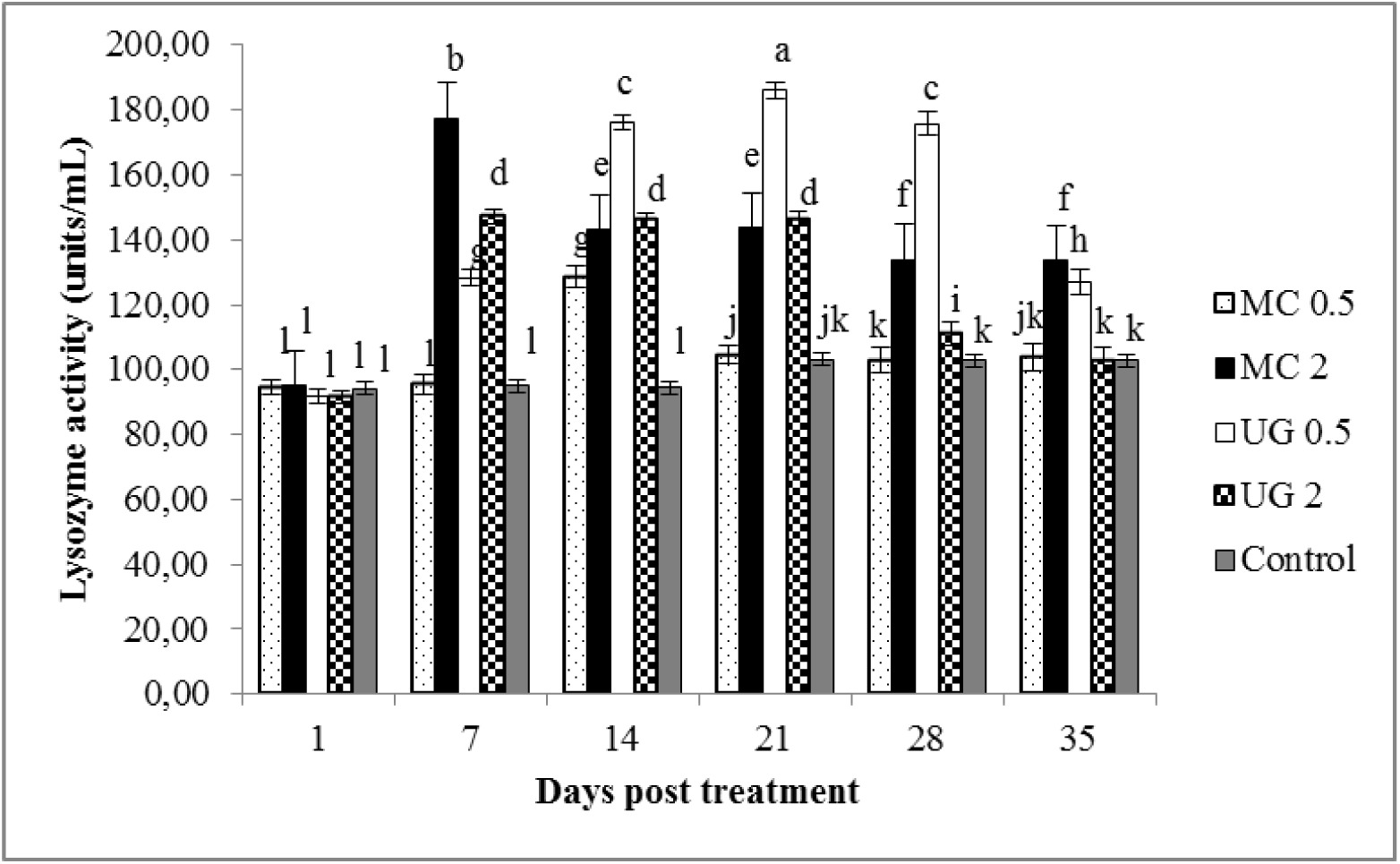
Effects of MC and UM extracts on the lysozyme activity in serum of European sea bass. Data are expressed as means ± SE (n = 10). Mean values at bars with different superscript letters at same stage are significantly different (P < 0.05) from the control.

### 3.3. Serum Total protein

The effect of plant extracts on the total protein in serum of fish is shown in Figure 3. There were no significant changes in serum total protein (P > 0.05).

**Figure 3.**
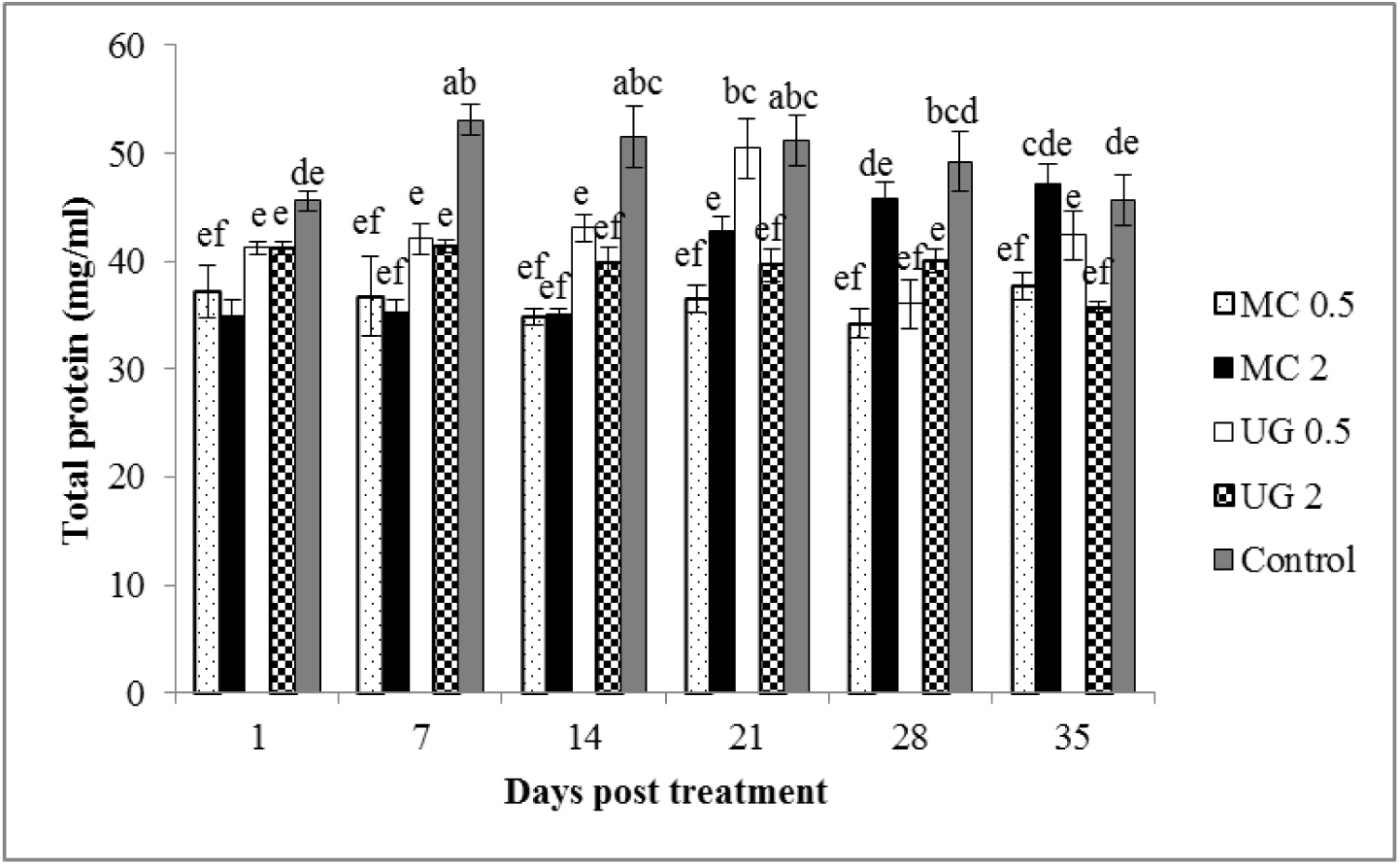
Effects of MC and UM extracts on the total protein level in serum of European sea bass. Data are expressed as means ± SE (n = 10). Mean values at bars with different superscript letters at same stage are significantly different (P > 0.05) from the control.

### 3.4. Hematology

The leukocyte numbers increased significantly (P < 0.05) in fish the injected with plant extracts. As seen in Table 1, fish showed elevated levels at both doses (0.5mg/fish MC or UM; 2mg/fish MC or UM) compared to the control group. The blood cell count of the fish injected with UM or MC plant extracts showed a significant increase in the number of monocytes and neutrophils (P< 0.05). The highest monocyte and neutrophil counts were observed in group 0.5mg/fish UM and 2mg/fish MC on day 21 (Table 1). the hematocrit levels in the samples did not show significant differences in any of the groups.

**Table 1.**
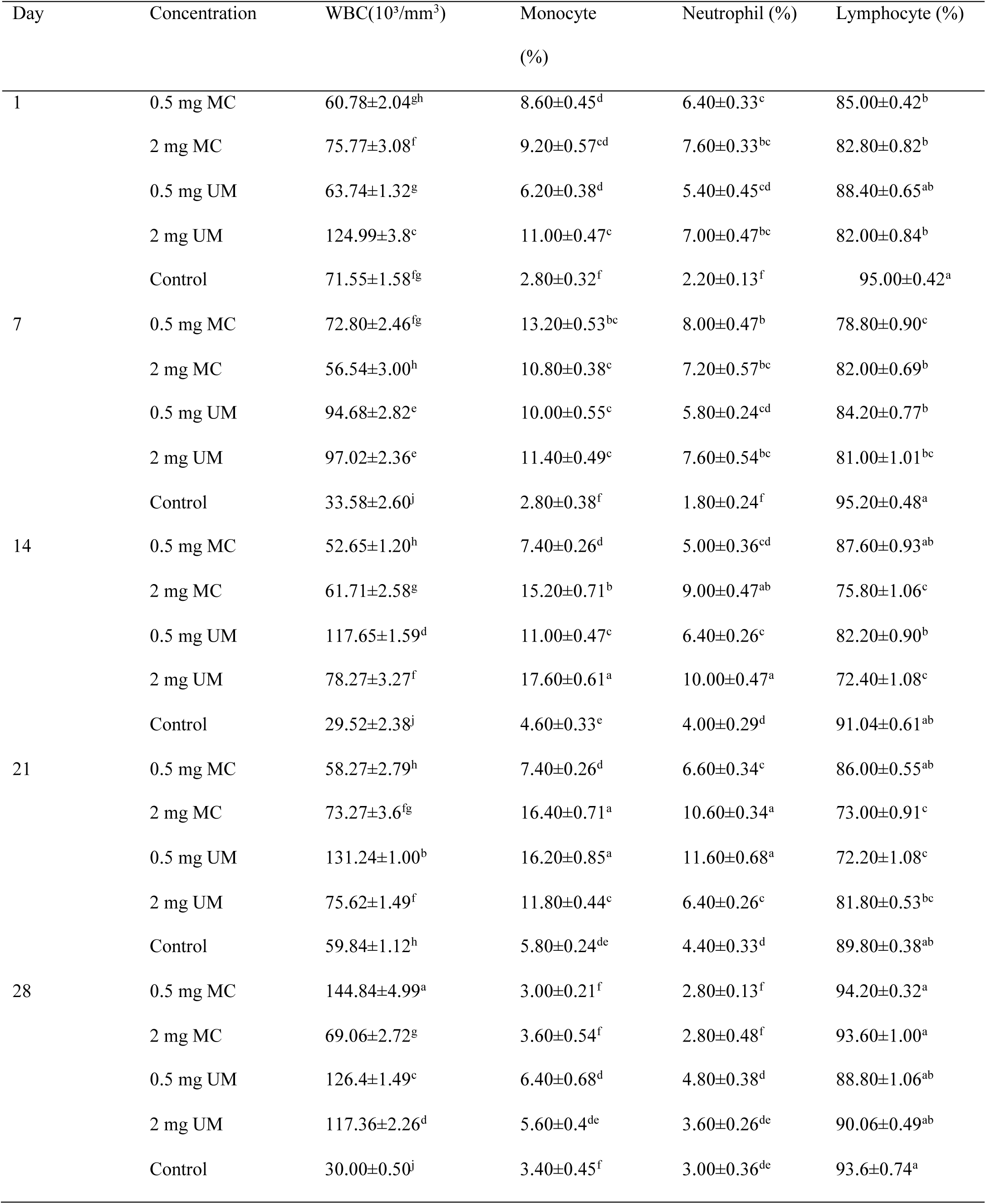
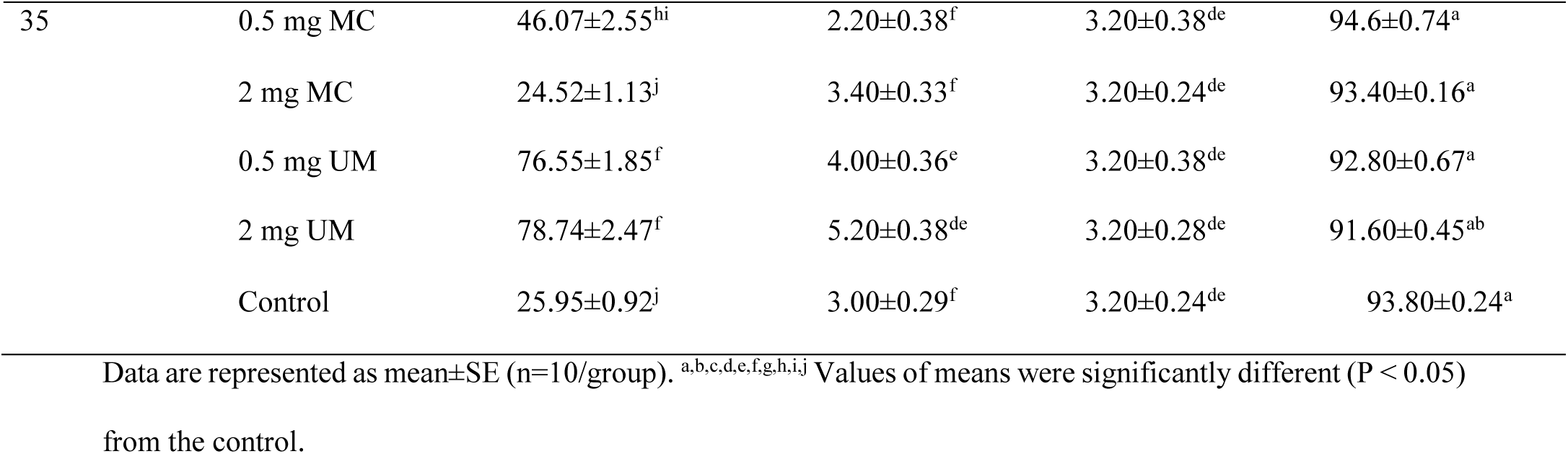
Effects of MC and UM extracts on differential blood count in European sea bass.

### 3.5. Growth

Specific growth rates (SGRs) of fish in each group were calculated and are presented in Table 2. Only the groups that received 0.5mg of MC or 2mg of MC per fish showed a significant difference (P > 0.05) in specific growth performance.

**Table 2.**
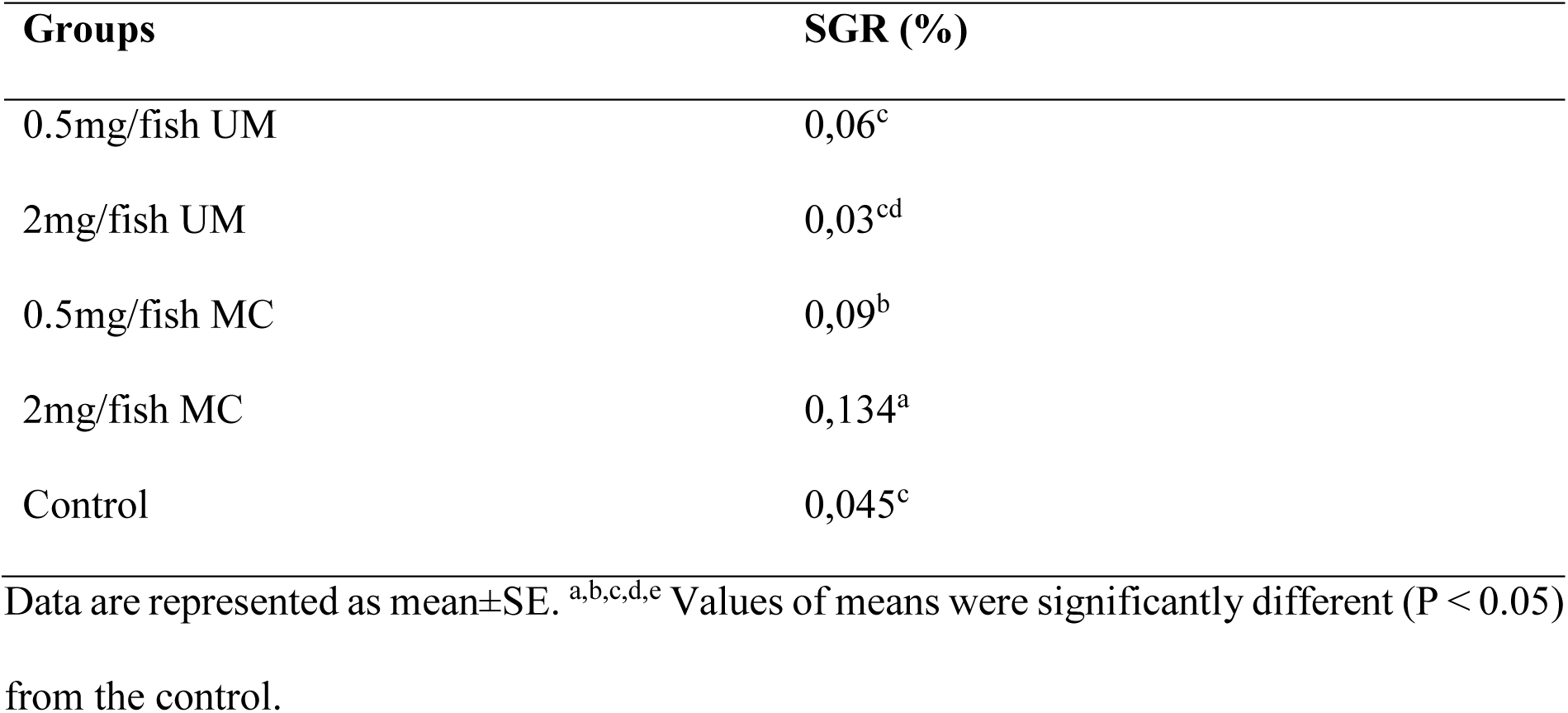
Effects of MC and UM extract on the Specific Growth Rate of European sea bass.

| Groups | SGR (%) |
| --- | --- |
| 0.5mg/fish UM | 0,06<sup>c</sup> |
| 2mg/fish UM | 0,03<sup>cd</sup> |
| 0.5mg/fish MC | 0,09<sup>b</sup> |
| 2mg/fish MC | 0,134<sup>a</sup> |
| Control | 0,045<sup>c</sup> |

Data are represented as mean±SE. <sup>a,b,c,d,e</sup> Values of means were significantly different (P < 0.05) from the control.

## 4. Discussion

Medicinal plants as immunostimulants in fin-fish sector have gotten more attention in the most recent decades, for their enhancing capacity of the immune system, as well as for their growth promoting effects (Sahu et al. 2007).

The present results show that fish injected with plant extracts significantly enhance non-specific immune parameters. Specifically, the application of UM and MC plant extracts in European sea bass (0.5 mg/fish and 2 mg/fish treatments) significantly increase the number of NBT positive phagocytic cells, achieving a maximum level on the 21th day in the 2 mg/fish MC experiment group. In comparative studies, in which fish were given immunostimulants by injection, it was reported that it took 2-4 days for the number of NBT-positive cells to increase (Anderson and Jeney 1992; Chen et al. 1998).

Comparative outcomes have been accounted for in various fish species, for example, *Oreochromis mossambicus*, (Logambal and Michael 2000), *Oncorhynchus mykiss* (Dügenci et al. 2003; Bilen et al. 2011), *Catla catla* (Rao and Chakrabarti 2005), *Carassius auratus* (Kumar et al. 2013), *Sparus aurata*, (Baba et al. 2014), *Oreochromis niloticus* (Zahran et al. 2014), *Victoria labeo* (Ngugi et al. 2015).

In the present study, European sea bass injected with MC and UM also exhibited increased lysozyme activity in serum. This is in agreement with earlier studies using plant derived substances. For example, by using 1% lupin *Lupinus perennis*, mango *Mangifera indica*, or stinging nettle *Urtica dioica* feeding rainbow trout with it for 14 days led to significant enhancement in serum lysozyme activity compared to the controls (Awad and Austin, 2010). In addition, rainbow trout that fed with plant extract such as 0.5% and 1% tetra *Cotinus coggyria* led to an increase in lysozyme activity compared to the control group; in particular, the 1% dose recorded the highest value (Bilen et al. 2011). Awad et al. (2013) recorded increases in lysozyme levels in rainbow trout fed with different doses of black cumin seed oil and nettle extract. Also Acar et al. (2015) showed increase in lysozyme activity in tilapia *O. mossambicus* fed with sweet orange *Citrus sinensis* essential oil. Previous reports, on lysozyme activity in fish which were treated with immunostimulants showed similar results to our study. (Hutchinson and Manning, 1996; Yeganeh et al. 2016)

Previous studies have examined the impact of plant extracts on total protein amounts in serum of fish. For example, in rainbow trout, increasing of plasma total protein level was found to be significant following the feeding of the fish with several herbal extracts (Dügenci et al. 2003). Binai et al. (2014) also recorded increases in total protein level in juvenile beluga fed with nettle. On the contrary, two Chinese medicinal herbs (*Astragalus membranaceus* and *Lonicera japonica*) and boron on non-specific immune response in Nile tilapia had no effect on plasma total protein (Ardo et al. 2008). In our study, serum total protein values did not change when compared to controls.

Hematocrit level is a indicator of general fish health and can indicate changes caused by immunostimulants (Mulero, 1998). In the current study, it was observed that the hematocrit level did not show any significant difference from the control group during the trial. Siwicki and Anderson (1993) demonstrated similar results for rainbow trout. Several studies in different fish species fed with *Spiruluna platensis* (Yeganeh et al. 2016) and *Citrus limon* (Ngugi et al. 2016) have reported similar results. In contrast to hematocrit, in the present study, the WBC levels in the blood of fish injected with UM and MC plant extracts significantly increased. The total white blood cell (WBC) count is an important indicator of immune response in fish (De Pedro et al., 2005). *Cyprinus carpio* injected intraperitoneally with β-glucan (100, 500 and 1000 mg) had a significant increase in total leukocyte counts and an increased proportion of neutrophils and monocytes on day 7 (Selvaraj et al. 2005). Also, many authors have documented an enhancement of the WBC level after using immunostimulants in fish (Kumar et al. 2013; Baba et al. 2015; Ngugi et al. 2016; Baba et al. 2016). Granulocytes have a vital role in non-specific defense in fish (Dalmo et al., 1997). In the present study, phagocytized peripheral blood leukocyte potential and killing activity of neutrophils and monocytes increased in UM and MC plant extracts injected fish. Similar findings have also been reported in some studies (Anderson and Jeney 1992, Awad and Austin 2010)

Several natural products have been tested previously for their growth promoting activity in aquaculture (Citarasu et al., 2002). This study shows that MC plant extracts improve the growth performance in European sea bass. By examining the specific growth rates, extracts applied at different doses do not have any negative effect on non-specific immune system, yet they have a positive effect on growth and improvement performance (Zahran et al 2014; Wang et al. 2015; Baba et al. 2016). The results of our study showed that the MC and UM plant extracts significantly enhance some non-specific immune parameters in European sea bass. Also the plant extracts have a positive effect on growth performance. Thus, it can be concluded that both plant extracts can be used as an immunostimulant to enhance non-specific immune response in sea bass.

## Acknowledgments

The authors are grateful to fish farm (Fyord Marin A..) for providing the necessary facilities to carry out this trials. The study was supported by TUBITAK Carrier Project (104V126). We would like to thank Emre-Nukhet YAVUZER, Derya-Muhammed ÖZEN for helping us during the trials. We also thank Süleyman BABA for his support in this study. We appreciate Ashley Neal HAINES for her valuable suggestions and corrections of the manuscript.

